# IRNdb: The database of immunologically relevant non-coding RNAs

**DOI:** 10.1101/037911

**Authors:** Elena Denisenko, Daniel Ho, Ousman Tamgue, Mumin Ozturk, Harukazu Suzuki, Frank Brombacher, Reto Guler, Sebastian Schmeier

## Abstract

MicroRNAs (miRNAs), long non-coding RNAs (lncRNAs) and other functional non-coding RNAs (ncRNAs) have emerged as pivotal regulators involved in multiple biological processes. Recently, ncRNA control of gene expression has been identified as a critical regulatory mechanism in the immune system. Despite the great efforts made to discover and characterize ncRNAs, the functional role for most remains unknown. To facilitate discoveries in ncRNA regulation of immune system-related processes we developed the database of immunologically relevant ncRNAs and target genes (IRNdb). We integrated mouse data on predicted and experimentally supported ncRNA-target interactions, ncRNA and gene annotations, biological pathways and processes, and experimental data in a uniform format with a user-friendly web interface. The current version of IRNdb documents 12,930 experimentally supported miRNA-target interactions between 724 miRNAs and 2,427 immune-related murine targets. In addition, we recorded 22,453 lncRNA-immune target and 377 PIWI-interacting RNA-immune target interactions. IRNdb is a comprehensive searchable data repository which will be of help in studying the role of ncRNAs in the immune system.

**Database URL:** http://irndb.org

## Introduction

Rapid development of high-throughput technologies enabled identification of non-coding RNAs (ncRNAs) as a highly abundant class of transcripts in the pervasively transcribed eukaryotic genome (1). Among ncRNAs, microRNAs (miRNAs), long non-coding RNAs (lncRNAs), and PIWI-interacting RNAs (piRNAs) have been attracting an increasing interest over the last decade as master regulators of numerous genes and diverse biological processes.

MiRNAs constitute a family of ~22 nucleotides small ncRNAs that bind to target mRNAs to mediate post-transcriptional repression or degradation of the mRNA (2). More than 20,000 miRNA loci are described to date in different species, including over 2,000 miRNAs in mammals (3). More than 50% of the mammalian genome is thought to be under miRNA control (4). Recently, a number of miRNAs were found to be important regulators acting in both adaptive and innate immune cells (5). Cooperative action of multiple miRNAs contributes to a systemic regulation of development and homeostasis of the immune system and the host response to invading pathogens (6–8). However, for the majority of miRNAs their roles in immune-related processes still remain to be elucidated.

PiRNAs constitute another class of small ncRNAs whose well-established function is to silence transposable elements in complex with PIWI proteins in germline cells (9). Recent studies provided evidence that piRNA might possess a wider range of functions including protein-coding gene silencing (including immune system-related genes) and epigenetic regulation (10).

LncRNAs are defined as ncRNAs longer than 200 nucleotides (11). Several classification systems and numerous subclasses of lncRNA genes have been described (12). LncRNAs are believed to possess a wide range of molecular functions including transcriptional regulation, regulation of mRNA processing, control of post-transcriptional events and regulation of protein activity (13). A number of lncRNAs were shown to regulate the expression of their adjacent immune genes and be functionally important in both innate and adaptive immunity (14). LncRNAs synthesized by both host and pathogen are involved in the regulation of host-pathogen interactions and might be crucial for the infection outcome (14). Despite the clear importance of certain lncRNAs for regulatory mechanisms, the functionality and biological role of the vast majority of lncRNAs remains unknown (13).

The numbers of experimental studies on ncRNA-target interactions as well as the numbers of computational prediction approaches and tools have increased drastically over the years (15,16). Accordingly, many databases have been developed, providing information on different aspects of ncRNA biology (17,18). Among these, miRBase serves as a central repository for up-to-date miRNA annotation and sequences (3). Experimentally validated ncRNA-target interactions are collected by several curated resources including miRTarBase (19), miRecords (20), LncRNA2Target (21), and piRBase (22). Different approaches were implemented for the computational prediction of miRNA-target interactions including TargetScan (23), microRNA.org (24), PITA (25), PicTar (26), MicroCosm (27) and many others. These tools often use a combination of prediction algorithms based on seed sequence match, free energy of miRNA-mRNA duplex, evolutionary conservation of the miRNA binding sequence, and other features of miRNA-target interactions (28). Specific resources including miR2Disease (29) and LncRNADisease (30) were developed to decipher the role of ncRNA in diseases.

In this study, we report the development of IRNdb, the first, to our knowledge, specialized database for immune-related ncRNA in mice, a model organism for the study of the immune system. IRNdb was developed with a goal of establishing a universal resource for murine ncRNAs in immunological research. We integrated information on murine miRNAs, lncRNAs, piRNAs, and their immunologically relevant target genes. We combined multiple sources of experimentally supported and predicted miRNA-target interactions, thus eliminating the user’s need to browse multiple websites and run different computational tools. In IRNdb we visualize information on experimentally supported and predicted interactions separately, thus, allowing users to choose a category of interest. NcRNAs and target genes are linked to external databases to offer additional information such as sequences and tissue-specific expression. We implemented a feature that enables users to perform functional analysis and identification of significant biological pathways for any selected subset of target genes using Enrichr, a popular tool which offers a large collection of gene set libraries (31). In addition to the identification of significant pathways, IRNdb provides a full list of biological pathways associated with the features of interest. This approach is particularly beneficial for short lists of targets, where over-representation analyses might fail to identify statistically significant pathways. In addition, we list transcription factor binding sites (TFBS) identified upstream of miRNA genes to study their transcriptional regulation. IRNdb has a simple user-friendly interface, is easily accessible and fully searchable, and provides functionality to download any findings. Hence, IRNdb offers a new data repository to improve our understanding of the roles of ncRNAs in the immune system.

## Data sources, integration and implementation

### Biological entities and interactions

We integrated public domain data from various sources (see Figure 1). Experimentally supported interactions between miRNAs and their target genes were retrieved from two databases: miRTarBase (19) and miRecords (20). Predicted interactions were obtained from nine tools/databases: DIANA microT-CDS (32), ElMMo3 (33), MicroCosm (27), microRNA.org (24), miRDB (34), miRNAMap2 (35), PicTar2 (DoRiNA 2.0) (26), PITA (25), and TargetScanMouse (23). We downloaded lists of miRNA-target predictions from the respective tool/database websites. These tools employ different prediction algorithms including seed sequence match, free energy of microRNA-mRNA duplex, and evolutionary conservation. We collected predicted miRNA-target interactions derived by at least two different tools. Experimentally supported and predicted miRNA-target interactions are listed separately in different tables; all corresponding sources are specified for each target. We only retained miRNAs and their interactions from the interactions resources where either a current miRBase ID was readily available or where the resource miRBase ID could be unambigously mapped to a miRBase accession through an alias/outdated miRBase IDs from miRBase. We also included lncRNA-target interaction from LncRNADisease (30), LncRNA2Target (21), and LncReg (36) and piRNA-target interactions from piRBase (22).

**Figure 1.**
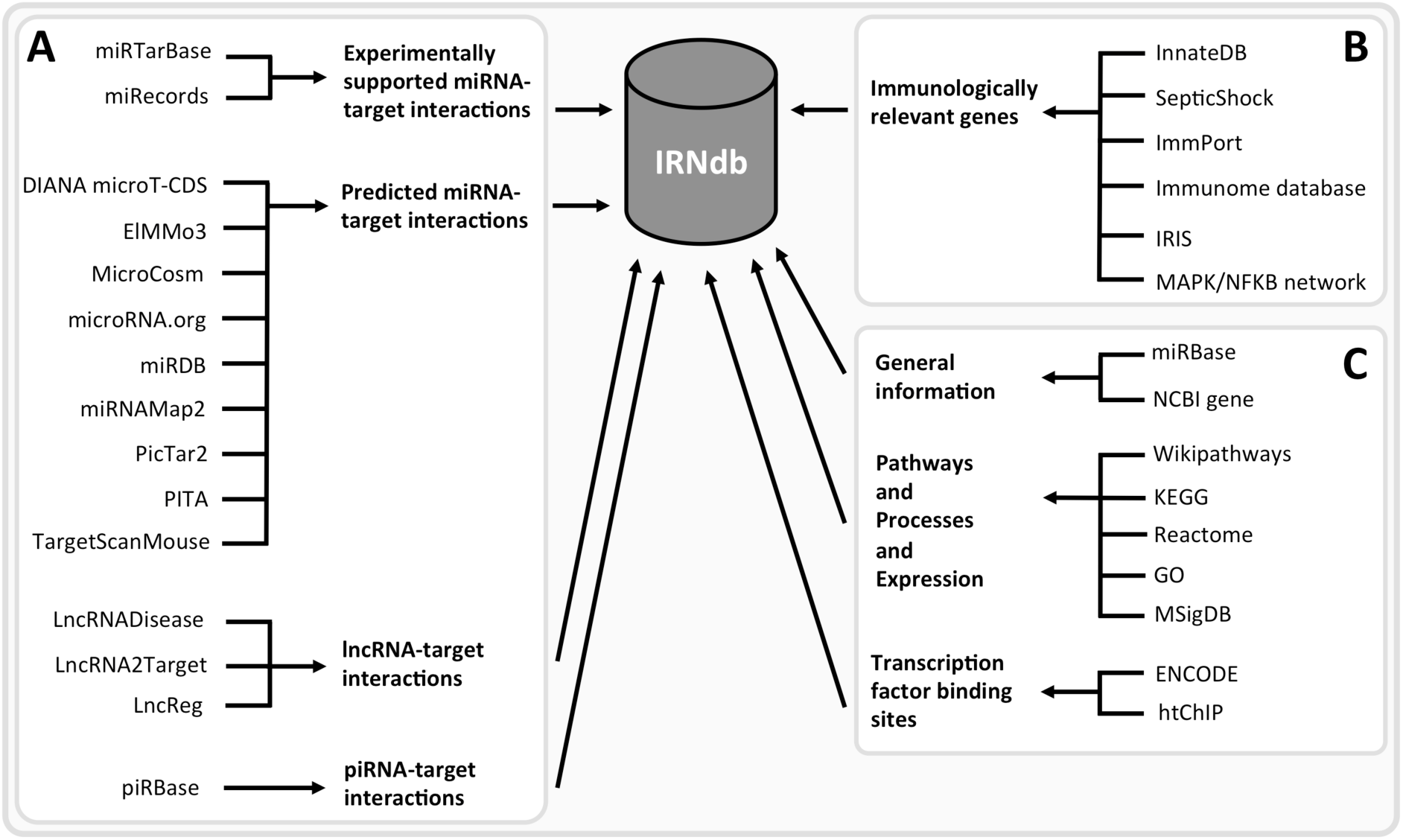
IRNdb construction: public domain data sources. (**A**) ncRNA-related information. (**B**) Information regarding immunologically relevant target genes. (**C**) Annotation data, e.g. general information regarding the biological entities, biological pathways, processes, gene expression data, transcription factor binding sites, etc.

The set of genes involved in immunological processes was retrieved from the following sources: InnateDB (37) and SepticShock (http://www.septicshock.org/) were used as sources of mouse-specific immunological gene information. Immunologically relevant human genes were extracted from ImmPort (38), the Immunome database (39,40), Immunogenetic Related Information Source (IRIS) (41), and InnateDB and the MAPK/NFKB network (37) and converted to murine genes via the NCBI Homologene database (http://www.ncbi.nlm.nih.gov/homologene). To date, IRNdb is focused solely on mouse data.

We retained ncRNAs in IRNdb if they could be connected to one of the immunologically relevant target genes through the above mentioned interaction databases. In total, we were able to extract 12,930 experimentally supported interactions between 724 miRNAs and 2,427 target genes (see Table 1). In addition, we collected 183,752 predicted interactions between 1,115 miRNAs and 5,297 targets (see Table 1). We only retained interactions predicted by at least two sources. However, many interactions were predicted through multiple databases/tools (see Table 2), e.g. over 42% of interactions were predicted by more than two sources and 0.22% of miRNA-gene interactions were predicted by all nine databases/tools. A miRNA interacts on average with 17.9 (164.8) target genes through experimentally supported (predicted) interactions. On the other hand, a target gene interacts on average with 5.3 (34.7) miRNAs through experimentally supported (predicted) interactions. In addition to miRNAs, we collected lncRNAs and piRNAs. We were able to extract 22,453 interactions between 163 lncRNAs and 4,521 genes and 377 interactions between 319 piRNAs and 84 genes. On average a lncRNA (piRNA) is connected to 137.7 (1.2) genes and a target gene is on average connected to 5 (4.5) lncRNAs (piRNAs).

**Table 1.** IRNdb data repository statistics on ncRNA-target gene interactions. If no version number was available we added the date of the last resource update we used.

|  | Immunological target information inferred from |  |
| --- | --- | --- |
|  | Mouse | Mouse+Human |
| **Experimentally supported microRNA-target interactions** | **4,451** | **12,930** |
| *miRTarBase* (v6.1) | 4,409 (99.1%) | 12,845 (99.3%) |
| *miRecords* (v5.0) | 118 (2.7%) | 245 (1.9%) |
| **Predicted microRNA-target interactions** | **60,824** | **183,752** |
| *DIANA microT-CDS* (v5) | 36,690 (60.3%) | 107,269 (58.4%) |
| *EIMMo3* (v3) | 6,443 (10.6%) | 18,476 (10.1%) |
| *MicroCosm* (v5) | 14,913 (24.5%) | 46,800 (25.5%) |
| *microRNA.org* (August 2010) | 26,986 (44.4%) | 80,566 (43.8%) |
| *miRDB* (v5) | 24,589 (40.4%) | 72,823 (39.6%) |
| *miRNAmap2* (v2.0) | 13,993 (23.0%) | 43,259 (23.5%) |
| *PicTar2* (v2) | 22,952 (37.7%) | 67,833 (36.9%) |
| *PITA* (v6) | 16,185 (26.6%) | 46,717 (25.4%) |
| *TargetScanMouse* (v7.1) | 16,450 (27.1%) | 48,219 (26.2%) |
| **Long non-coding RNA-target interactions** | **7,778** | **22,453** |
| *LncRNADisease* (June 2015) | 13 (0.2%) | 29 (0.1%) |
| *LncRNA2Target* (v1.2) | 7,669 (98.6%) | 22,250 (99.1%) |
| *LncReg* (2015) | 105 (1.4%) | 190 (0.9%) |
| **PIWI-interacting RNA-target interactions** | **141** | **377** |
| *piRBase* (v1.0) | 141 (100%) | 377 (100%) |

**Table 2.** Percentage of predicted microRNA-target interactions covered by multiple sources.

|  | **Immunological target information**<br>**inferred from** |  |
| --- | --- | --- |
|  | **Mouse** | **Mouse+Human** |
| **Predicted**<br>**microRNA-target**<br>**interactions** | **60,824** | **183,752** |
| **Percentage of**<br>**interactions covered by**<br>**this many of sources** | **%** | **%** |
| 9 | 0.23 | 0.22 |
| 8 | 0.97 | 0.86 |
| 7 | 2.62 | 2.35 |
| 6 | 4.16 | 3.71 |
| 5 | 5.66 | 5.49 |
| 4 | 9.96 | 9.73 |
| 3 | 20.60 | 20.30 |
| 2 | 55.80 | 57.34 |

### Biological annotation data

General information on miRNAs and corresponding targets was extracted from miRBase (3) and NCBI gene (https://www.ncbi.nlm.nih.gov/gene). Biological pathway and process information was downloaded from Kyoto Encyclopedia of Genes and Genomes (KEGG) (42), WikiPathways (43), Gene Ontology (GO) (44), and Reactome (45).

In addition, lists of up- and down-regulated genes in immunologically relevant experiments were downloaded from the Molecular Signature Database (MSigDB) (46). We integrated information on relative expression of 455 mature miRNAs in 68 small RNA libraries (47) to indicate mouse cells and tissues where miRNAs were shown to be expressed.

Transcription factor binding site (TFBS) information was retrieved from ENCODE (48) and HT-ChIP (49). We mapped raw sequencing data to the mm10 genome build for each tissue and cell type separately and called peaks using MACS2 (50). We retained TFBS peaks within 2500bp upstream of the 5’end of primary miRNAs with a threshold of FDR < 0.01. For each mature miRNA and all corresponding precursors we record the identified transcription factors (TF), distance between TFBS and miRNA gene, corresponding FDR and cell type.

### Database implementation

IRNdb (accessible at http://irndb.org) is an open-access database implemented in the DJANGO web-framework (https://www.djangoproject.com/), running on an NGINX web-server (https://www.nginx.com/) with a MySQL database server (https://www.mysql.com/) for storing data and Memcached (https://memcached.org/) for object caching. The web-frontend is developed using the Bootstrap framework (http://getbootstrap.com/). The source code for IRNdb is freely available at https://github.com/sschmeier/irndb2.

## Website usage

### Database and web-interface overview

IRNdb collects information on ncRNAs targeting immunologically relevant protein-coding genes and biological annotation data (see Figure 1). To access the content in IRNdb, we developed a web-interface (accessible at http://irndb.org), which is divided into four main sections ‘microRNAs’, ‘piRNAs’, ‘lncRNAs’, and ‘Target genes’. In addition, the ‘Documentation’ section provides instructions and examples for using the database, contact information, and a statistical summary of the IRNdb interaction data. Below we illustrate IRNdb features and functionality using miRNAs mmu-miR-125b-5p and mmu-miR-873a-5p, as well as their targets as examples.

### The ncRNA-centred view

IRNdb provides an interface for the convenient retrieval of ncRNA and target information. Users can browse ncRNAs in tabular form that is searchable, e.g. by identifier or names. Clicking the ncRNA name in the table leads to an individual ncRNA entry (e.g. ‘miRNA view’ in case of miRNAs).

For illustrative purposes, we focus on the role of mmu-miR-125b-5p in mouse macrophages. One of the known functions of miR-125b is targeting TNF-alpha mRNA in macrophages in response to mycobacterial infection (51). An individual ‘miRNA view’ for mmu-miR-125b-5p provides five tabs with information regarding the ncRNA (see Figure 2A). The ‘Targets’-tab contains separate tables for experimentally supported and predicted ncRNA targets. In IRNdb we visualize information on experimentally supported and predicted interactions separately, thus, allowing users to choose a category of interest. We indicate sources of every interaction so that users can use all of them or focus on the most reliable interactions predicted by many tools. The tables are searchable and Tnf can be found among experimentally supported targets. Searching for ‘Tnf’ among predicted targets reveals that Tnf receptor, tumor necrosis factor receptor superfamily member 1b (Tnfrsf1b), is predicted to be a target of mmu-miR-125b-5p. The column ‘Target source’ indicates three different sources for this prediction. To evaluate this potential target, we can switch to IRNdb ‘Target gene view’ by clicking the gene symbol link (see Figure 2A). The ‘Expression’ field of the ‘Target gene view’ provides three links to different external sources where we can check if Tnfrsf1b is expressed in macrophages (see Figure 2B). The first link opens the EBI Expression Atlas (52) entry for murine Tnfrsf1b. Under the EBI Expression Atlas ‘Cell type’ header we find that Tnfrsf1b has a medium expression level in mouse macrophages. The second link on the IRNdb target view opens the FANTOM5 SSTAR catalog entry for Tnfrsf1b (http://fantom.gsc.riken.jp/5/sstar) which, again, shows that Tnfrsf1b is expressed in mouse macrophages. Finally, we provide a link to TBvis, a resource where we implemented visualization of cap analysis gene expression (CAGE) (53) data for murine macrophages cultivated under different conditions and infected with *Mycobacterium tuberculosis* (*Mtb*) (54,55). TBvis data for Tnf and Tnfrsf1b indicate that both genes show an increase in their expression in macrophages upon infection with *Mtb*.

**Figure 2.**
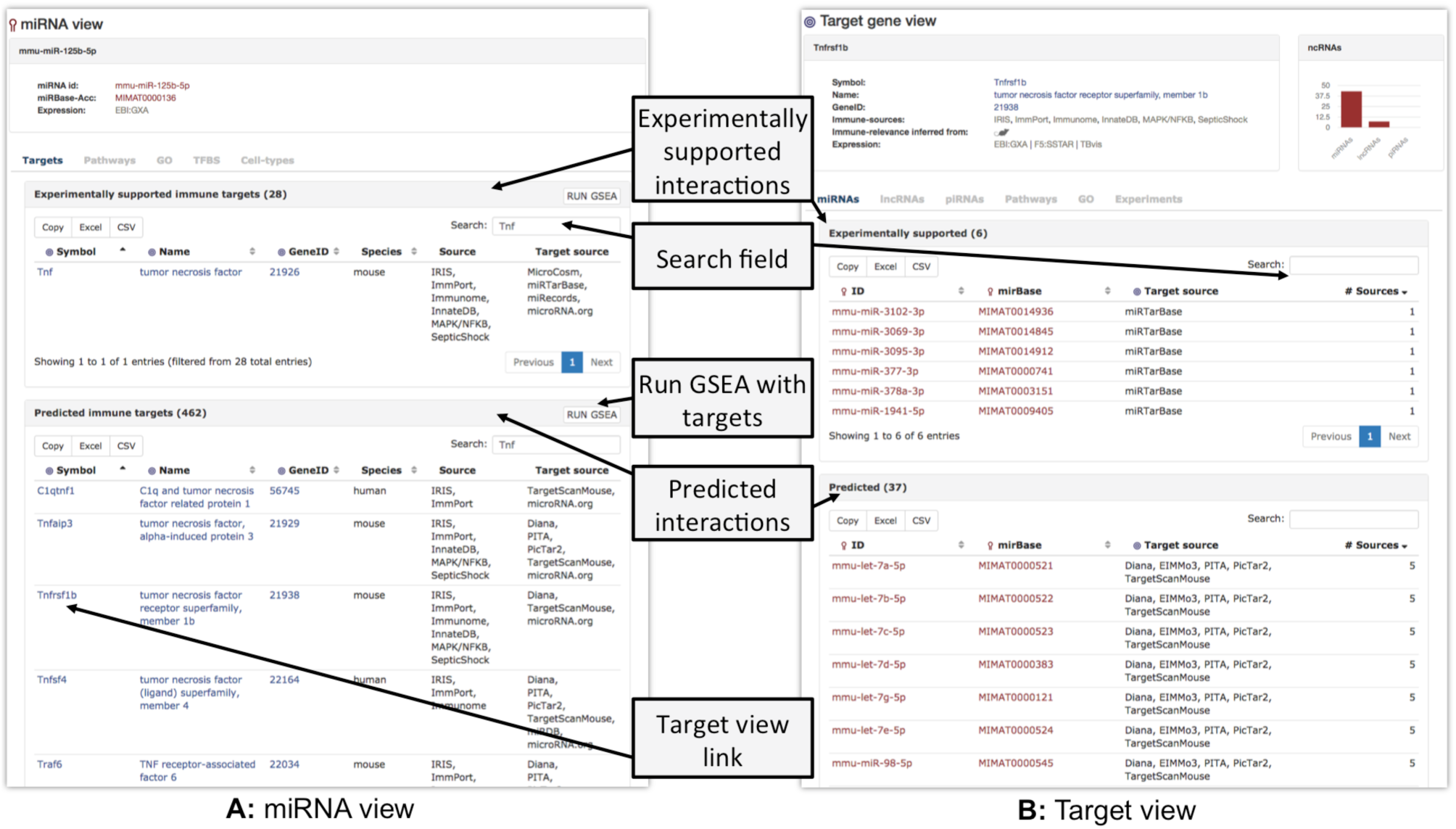
Overview of the ‘ncRNA’-view and ‘Target gene’-view.

If regulation of miRNA transcription is of interest, users can browse the table in the ‘TFBS’-tab of the ‘miRNA view’. The table lists transcription factors for which TFBSs were found upstream of miRNA in question, along with additional information on cell type, TFBS location and significance. The ‘Cell types’-tab shows relative cloning frequencies of corresponding miRNA across a set of murine tissues and cell types (47). Macrophages were not profiled in this experiment; however, we find that mmu-miR-125b-5p is expressed in other immune cell types including T-, B-, and dendritic cells, which might indicate a complex role of mmu-miR-125b-5p in response to the infection.

### Browsing and searching target genes

IRNdb provides a separate search interface for immune system-related ncRNA target genes. Genes can be searched using gene symbol, gene name, Entrez Gene ID, or their parts. Moreover, users can select any set of genes by clicking on the corresponding rows and perform the gene set enrichment analysis (GSEA) by Enrichr tool (31) with ‘RUN GSEA’ button. The table provides information on the numbers of experimentally supported and predicted miRNAs, lncRNAs, and piRNAs that target each gene. Users can switch to the ‘Target gene view’ while browsing ncRNA targets by clicking the gene symbol link.

### The target-centred view

We continue the demonstration of how IRNdb can be used to characterize Tnfrsf1b, the predicted target of mmu-miR-125b-5p. In addition to the aforementioned ‘Expression’ field with three links to the external sources of tissue and cell type-specific gene expression, the ‘Target gene view’ provides six tabs (see Figure 2B).

The ‘miRNAs’, ‘lncRNAs’, and ‘piRNAs’-tabs show tables with corresponding ncRNAs that target the gene in question. To date, for Tnfrsf1b IRNdb lists seven lncRNAs, six experimentally supported miRNAs, and 37 predicted miRNAs. In the case of miRNAs, each table can be sorted by the number of sources of miRNA-target information. Thus, if one aims at testing some of the predicted Tnfrsf1b-targeting miRNAs, the mmu-let-7 family and mmu-miR-98-5p might represent miRNAs of interest as they were predicted by the largest number of tools (see Data sources, integration and implementation).

The ‘Pathways’- and ‘GO’-tabs provide pathways and processes associated with the target gene. They indicate that Tnfrsf1b is involved in apoptosis and oxidative damage-related processes. The ‘Experiments’-tab provides additional functional information on the target, listing immunologically relevant experiments where the target was shown to be up- or down-regulated. Searching for ‘macrophages’ reveals that Tnfrsf1b was found to be up- or down-regulated in multiple experiments on macrophages. The experiments can be searched by any key words to retrieve, for instance, disease, stimulation, or cell type of interest. We integrated experiments performed for mouse and human which is indicated in the “Species” field.

### Pathway analysis of ncRNA targets

We implemented two complementary approaches for ncRNA functional analysis via analysis of the corresponding target genes. We demonstrate how IRNdb can be used for functional analysis of ncRNAs and their targets on the example of miRNA mmu-miR-873a-5p.

A list of targets of a ncRNA of interest can be subjected to the gene set enrichment analysis (GSEA) by Enrichr tool (31) by pressing the ‘RUN GSEA’ button of the ‘Targets’-tab of the ncRNA view. In the case of miRNAs, experimentally supported and predicted targets are analysed separately. Pressing the ‘RUN GSEA’ button for predicted targets of mmu-miR-873a-5p launches Enrichr in a new window, where users can browse numerous different biological categories. Among KEGG and WikiPathways terms, predicted targets of mmu-miR-873a-5p were significantly enriched in B cell receptor, IL-4, and TNF signaling pathways.

The list of experimentally supported targets can be analysed in a similar manner. However, in the case of mmu-miR-873a-5p, only six experimentally supported targets are reported in IRNdb to date. In such case, it might be beneficial to browse all pathways associated with these targets instead of focusing on statistically significant ones, as with such a small target set, no significance can be reliably calculated. With this purpose we created ‘Pathways’-tab of ncRNA view. It allows users to browse WikiPathways, KEGG, and Reactome pathways that contain at least one target of the ncRNA under investigation. Pathways are presented as a table for each pathway source respectively. Using this feature we found that one of mmu-miR-873a-5p experimentally supported targets, tumor necrosis factor, alpha-induced protein 3 (Tnfaip3), is associated with the WikiPathway entry TNF-alpha NF-kB Signaling Pathway. These data combined propose that mmu-miR-873a-5p might have a strong role in Tnf signaling which is in agreement with previously published data (56). Moreover, mmu-miR-873a-5p can be further studied to elucidate its possible involvement in the regulation of B cell receptor and IL-4 signaling pathways.

Similarly to the ‘Pathways’-tab, the ‘GO’-tab provides users with a list of GO-process and GO-function terms associated with the ncRNA targets. We believe that exploring the ‘GO’- and ‘Pathways’-tab, in addition to GSEA, enhances IRNdb capability of identification of biological pathways and processes possibly affected by the ncRNA of interest.

### Biological pathways

The IRNdb repository includes information on biological pathways with immunologically relevant ncRNA target genes. Users can reach the pathway data in three different ways by selecting the pathway from the ncRNA view’s ‘Pathways’-tab, from the target view’s ‘Pathways’-tab, or from the ‘by pathways’-option for a ncRNA class of interest in the navigation panel. In the latter case, the link leads to a ‘Browse pathways’-page, where IRNdb lists for each pathway the immunologically relevant experimentally supported ncRNA target genes and corresponding targeting ncRNAs.

Here, we focus on ‘TNF-alpha NF-kB Signaling Pathway’ of the WikiPathways catalogue. As discussed above, two miRNAs mmu-miR-125b-5p and mmu-miR-873a-5p could play a role in regulating this pathway. After searching for the pathway in the table on the ‘Wikipathway’-tab, we enter a pathway-centric view by clicking on the pathway name. The ‘miRNAs’-tab of the ‘Pathway view’ lists miRNAs with experimentally supported or predicted targets in the pathway. Searching for ‘mmu-miR-873a-5p’ shows that, while Tnfaip3 is its only experimentally supported target in the pathway, mmu-miR-873a-5p is predicted to target an additional 13 genes in the pathway, including Ikbkb and Nfkbie which regulate the NF-kappa-B transcription factor (57). Interestingly, mmu-miR-873a-3p has two predicted targets in the pathway as well. Similarly, mmu-miR-125b-5p has three experimentally supported and an additional 18 predicted targets in the ‘TNF-alpha NF-kB Signaling Pathway’. ‘LncRNA’ and ‘piRNA’-tabs provide tables with corresponding ncRNAs and their pathway associated targets.

## Conclusion

Over the past years, research into ncRNAs has increased significantly within the scientific community. For example, several miRNA databases with diverse scopes and goals have been recently introduced (28,58–60). MirDIP (28) and miRGate (59) were developed to address the issue of poor overlap among target prediction tools. A group of resources aims at deciphering biological functions of miRNAs. DIANA-miRPath (61) implements enrichment analysis of miRNA targets in Kyoto Encyclopedia of Genes and Genomes (KEGG) pathways (42) and Gene Ontology (GO) terms (44). MiRGator combines three miRNA target prediction programs and focuses on the expression analysis and gene set enrichment analysis to facilitate functional annotation of miRNAs (62). MiTALOS accounts for miRNA tissue specificity by introducing a novel pathway analysis methodology (63).

Undoubtedly, these efforts lead to a better data accessibility and understanding of ncRNA biology and function. One of the insights in ncRNA control is that it is exerted in a tissue- and condition-specific manner. Hence, without a specialized resource, an investigation of ncRNA roles under a certain condition might require time-consuming navigation between multiple data sources and gathering scattered condition-specific information from published literature. To bridge the gap in the understanding of ncRNA control of the immune system we developed IRNdb. To our knowledge, this is the first resource to focus on immune-related murine ncRNAs and their target genes. IRNdb is an integrative, user-friendly and easy-to-use platform with a wide range of applications (see Figure 3). IRNdb brings together experimentally supported and predicted ncRNA-target interactions enriched with pathway, GO, and experimental information, which will benefit ncRNA research and will help in the functional characterization of ncRNAs. All data tables in IRNdb are searchable and downloadable in a variety of formats. In the future we aim at integrating information on additional types of ncRNAs and expand IRNdb to include more mammalian systems.

**Figure 3.**
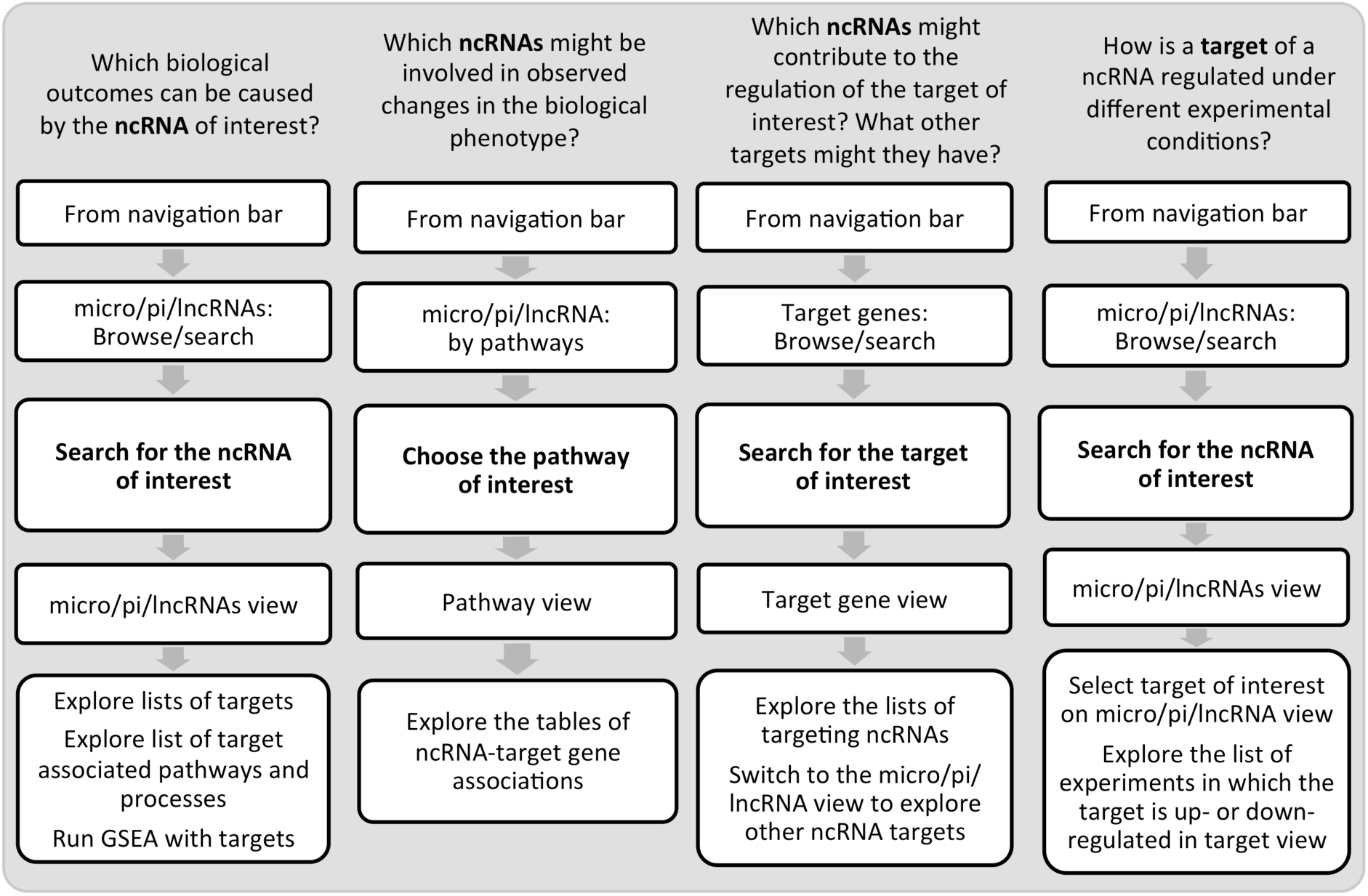
Examples of IRNdb applications in immunological research.

## Acknowledgements

This work was supported by the NRF Competitive Programme for Unrated Researchers (CSUR) to RG and SS.

## References

1. Sabin, L.R., Delas, M.J. and Hannon, G.J. (2013) Dogma derailed: the many influences of RNA on the genome. Mol. Cell, 49, 783–794.

2. Bartel, D.P. (2004) MicroRNAs: genomics, biogenesis, mechanism, and function. Cell, 116, 281–297.

3. Kozomara, A. and Griffiths-Jones, S. (2014) miRBase: annotating high confidence microRNAs using deep sequencing data. Nucleic Acids Res., 42, D68–73.

4. Friedman, R.C., Farh, K.K.-H., Burge, C.B. et al. (2009) Most mammalian mRNAs are conserved targets of microRNAs. Genome Res., 19, 92–105.

5. Gracias, D.T. and Katsikis, P.D. (2011) MicroRNAs: key components of immune regulation. Adv. Exp. Med. Biol., 780, 15–26.

6. Xiao, C. and Rajewsky, K. (2009) MicroRNA control in the immune system: basic principles. Cell, 136, 26–36.

7. Guler, R. and Brombacher, F. (2015) Host-directed drug therapy for tuberculosis. Nat. Chem. Biol., 11, 748–751.

8. Schmeier, S., MacPherson, C.R., Essack, M. et al. (2009) Deciphering the transcriptional circuitry of microRNA genes expressed during human monocytic differentiation. BMC Genomics, 10, 595.

9. Siomi, M.C., Sato, K., Pezic, D. et al. (2011) PIWI-interacting small RNAs: the vanguard of genome defence. Nat. Rev. Mol. Cell Biol., 12, 246–258.

10. Ku, H.-Y. and Lin, H. (2014) PIWI proteins and their interactors in piRNA biogenesis, germline development and gene expression. National Science Review, 1, 205–218.

11. Kung, J.T., Colognori, D. and Lee, J.T. (2013) Long noncoding RNAs: past, present, and future. Genetics, 193, 651–669.

12. St Laurent, G., Wahlestedt, C. and Kapranov, P. (2015) The Landscape of long noncoding RNA classification. Trends Genet., 10.1016/j.tig.2015.03.007

13. Geisler, S. and Coller, J. (2013) RNA in unexpected places: long non-coding RNA functions in diverse cellular contexts. Nat. Rev. Mol. Cell Biol., 14, 699–712.

14. Heward, J.A. and Lindsay, M.A. (2014) Long non-coding RNAs in the regulation of the immune response. Trends Immunol., 35, 408–419.

15. Leitao, A.L., Costa, M.C. and Enguita, F.J. (2014) A guide for miRNA target prediction and analysis using web-based applications. Methods Mol. Biol. (N. Y., NY, U. S.), 1182, 265–277.

16. Thomas, M., Lieberman, J. and Lal, A. (2010) Desperately seeking microRNA targets. Nat. Struct. Mol. Biol., 17, 1169–1174.

17. Fritah, S., Niclou, S.P. and Azuaje, F. (2014) Databases for lncRNAs: a comparative evaluation of emerging tools. RNA, 20, 1655–1665.

18. Akhtar, M.M., Micolucci, L., Islam, M.S. et al. (2015) Bioinformatic tools for microRNA dissection. Nucleic Acids Res., 10.1093/nar/gkv1221

19. Hsu, S.D., Tseng, Y.T., Shrestha, S. et al. (2014) miRTarBase update 2014: an information resource for experimentally validated miRNA-target interactions. Nucleic Acids Res., 42, D78–85.

20. Xiao, F., Zuo, Z., Cai, G. et al. (2009) miRecords: an integrated resource for microRNA-target interactions. Nucleic Acids Res., 37, D105–110.

21. Jiang, Q., Wang, J., Wu, X. et al. (2015) LncRNA2Target: a database for differentially expressed genes after lncRNA knockdown or overexpression. Nucleic Acids Res., 43, D193–196.

22. Zhang, P., Si, X., Skogerbo, G. et al. (2014) piRBase: a web resource assisting piRNA functional study. Database, 2014, bau110.

23. Grimson, A., Farh, K.K., Johnston, W.K. et al. (2007) MicroRNA targeting specificity in mammals: determinants beyond seed pairing. Mol. Cell, 27, 91–105.

24. Betel, D., Wilson, M., Gabow, A. et al. (2008) The microRNA.org resource: targets and expression. Nucleic Acids Res., 36, D149–153.

25. Kertesz, M., Iovino, N., Unnerstall, U. et al. (2007) The role of site accessibility in microRNA target recognition. Nat. Genet., 39, 1278–1284.

26. Blin, K., Dieterich, C., Wurmus, R. et al. (2015) DoRiNA 2.0--upgrading the doRiNA database of RNA interactions in post-transcriptional regulation. Nucleic Acids Res., 43, D160–167.

27. Enright, A.J., John, B., Gaul, U. et al. (2003) MicroRNA targets in Drosophila. Genome Biol., 5, R1.

28. Shirdel, E.A., Xie, W., Mak, T.W. et al. (2011) NAViGaTing the Micronome – Using Multiple MicroRNA Prediction Databases to Identify Signalling Pathway-Associated MicroRNAs. PLoS One, 6, e17429.

29. Jiang, Q., Wang, Y., Hao, Y. et al. (2009) miR2Disease: a manually curated database for microRNA deregulation in human disease. Nucleic Acids Res., 37, D98–104.

30. Chen, G., Wang, Z., Wang, D. et al. (2013) LncRNADisease: a database for long-non-coding RNA-associated diseases. Nucleic Acids Res., 41, D983–986.

31. Chen, E.Y., Tan, C.M., Kou, Y. et al. (2013) Enrichr: interactive and collaborative HTML5 gene list enrichment analysis tool. BMC Bioinf., 14, 128.

32. Paraskevopoulou, M.D., Georgakilas, G., Kostoulas, N. et al. (2013) DIANA-microT web server v5.0: service integration into miRNA functional analysis workflows. Nucleic Acids Res., 41, W169–173.

33. Gaidatzis, D., van Nimwegen, E., Hausser, J. et al. (2007) Inference of miRNA targets using evolutionary conservation and pathway analysis. BMC Bioinf., 8, 69.

34. Wong, N. and Wang, X. (2015) miRDB: an online resource for microRNA target prediction and functional annotations. Nucleic Acids Res., 43, D146–152.

35. Hsu, S.D., Chu, C.H., Tsou, A.P. et al. (2008) miRNAMap 2.0: genomic maps of microRNAs in metazoan genomes. Nucleic Acids Res., 36, D165–169.

36. Zhou, Z., Shen, Y., Khan, M.R. et al. (2015) LncReg: a reference resource for lncRNA-associated regulatory networks. Database, 10.1093/database/bav083

37. Breuer, K., Foroushani, A.K., Laird, M.R. et al. (2013) InnateDB: systems biology of innate immunity and beyond--recent updates and continuing curation. Nucleic Acids Res., 41, D1228–1233.

38. Bhattacharya, S., Andorf, S., Gomes, L. et al. (2014) ImmPort: disseminating data to the public for the future of immunology. Immunol. Res., 58, 234–239.

39. Ortutay, C., Siermala, M. and Vihinen, M. (2007) Molecular characterization of the immune system: emergence of proteins, processes, and domains. Immunogenetics, 59, 333–348.

40. Ortutay, C. and Vihinen, M. (2006) Immunome: a reference set of genes and proteins for systems biology of the human immune system. Cell. Immunol., 244, 87–89.

41. Kelley, J., de Bono, B. and Trowsdale, J. (2005) IRIS: a database surveying known human immune system genes. Genomics, 85, 503–511.

42. Kanehisa, M. and Goto, S. (2000) KEGG: kyoto encyclopedia of genes and genomes. Nucleic Acids Res., 28, 27–30.

43. Kelder, T., van Iersel, M.P., Hanspers, K. et al. (2012) WikiPathways: building research communities on biological pathways. Nucleic Acids Res., 40, D1301–1307.

44. Ashburner, M., Ball, C.A., Blake, J.A. et al. (2000) Gene ontology: tool for the unification of biology. The Gene Ontology Consortium. Nat. Genet., 25, 25–29.

45. Fabregat, A., Sidiropoulos, K., Garapati, P. et al. (2016) The Reactome pathway Knowledgebase. Nucleic Acids Res., 44, D481–487.

46. Liberzon, A., Subramanian, A., Pinchback, R. et al. (2011) Molecular signatures database (MSigDB) 3.0. Bioinformatics, 27, 1739–1740.

47. Landgraf, P., Rusu, M., Sheridan, R. et al. (2007) A Mammalian microRNA Expression Atlas Based on Small RNA Library Sequencing. Cell, 129, 1401–1414.

48. Yue, F., Cheng, Y., Breschi, A. et al. (2014) A comparative encyclopedia of DNA elements in the mouse genome. Nature, 515, 355–364.

49. Garber, M., Yosef, N., Goren, A. et al. (2012) A High-Throughput Chromatin Immunoprecipitation Approach Reveals Principles of Dynamic Gene Regulation in Mammals. Mol. Cell, 47, 810–822.

50. Zhang, Y., Liu, T., Meyer, C.A. et al. (2008) Model-based analysis of ChIP-Seq (MACS). Genome Biol., 9, R137.

51. Bettencourt, P., Pires, D. and Anes, E. (2016) Immunomodulating microRNAs of mycobacterial infections. Tuberculosis (Oxford, U. K.), 97, 1–7.

52. Petryszak, R., Burdett, T., Fiorelli, B. et al. (2014) Expression Atlas update--a database of gene and transcript expression from microarray- and sequencing-based functional genomics experiments. Nucleic Acids Res., 42, D926–932.

53. Kanamori-Katayama, M., Itoh, M., Kawaji, H. et al. (2011) Unamplified cap analysis of gene expression on a single-molecule sequencer. Genome Res., 21, 1150–1159.

54. Arner, E., Daub, C.O., Vitting-Seerup, K. et al. (2015) Gene regulation. Transcribed enhancers lead waves of coordinated transcription in transitioning mammalian cells. Science, 347, 1010–1014.

55. Roy, S., Schmeier, S., Arner, E. et al. (2015) Redefining the transcriptional regulatory dynamics of classically and alternatively activated macrophages by deepCAGE transcriptomics. Nucleic Acids Res., 43, 6969–6982.

56. Liu, X., He, F., Pang, R. et al. (2014) Interleukin-17 (IL-17)-induced microRNA 873 (miR-873) contributes to the pathogenesis of experimental autoimmune encephalomyelitis by targeting A20 ubiquitin-editing enzyme. J. Biol. Chem., 289, 28971–28986.

57. Halsey, T.A., Yang, L., Walker, J.R. et al. (2007) A functional map of NFκB signaling identifies novel modulators and multiple system controls. Genome Biol., 8, 1–14.

58. Ulfenborg, B., Jurcevic, S., Lindlof, A. et al. (2015) miREC: a database of miRNAs involved in the development of endometrial cancer. BMC Res. Notes, 8, 104.

59. Andrés-León, E., González Peña, D., Gómez-López, G. et al. (2015) miRGate: a curated database of human, mouse and rat miRNA–mRNA targets. Database, 10.1093/database/bav035

60. Schmeier, S., Schaefer, U., MacPherson, C.R. et al. (2011) dPORE-miRNA: polymorphic regulation of microRNA genes. PLoS One, 6, e16657.

61. Vlachos, I.S., Zagganas, K., Paraskevopoulou, M.D. et al. (2015) DIANA-miRPath v3.0: deciphering microRNA function with experimental support. Nucleic Acids Res., 43, W460–466.

62. Nam, S., Kim, B., Shin, S. et al. (2008) miRGator: an integrated system for functional annotation of microRNAs. Nucleic Acids Res., 36, D159–164.

63. Preusse, M., Theis, F.J. and Mueller, N.S. (2016) miTALOS v2: Analyzing Tissue Specific microRNA Function. PLoS One, 11, e0151771.

